# High rates of human faecal carriage of *mcr-1-*positive multi-drug resistant isolates emerge in China in association with successful plasmid families

**DOI:** 10.1101/106575

**Authors:** Lan-Lan Zhong, Hang TT Phan, Xi Huang, Karina Doris-Vihta, Anna E Sheppard, Kun-Jiao Zeng, Hong-Yu Li, Xue-Fei Zhang, Sandip Patil, Yan-Fen Zhang, Cong Shen, Derrick W Crook, A Sarah Walker, Yong Xing, Qian-yi Chen, Jia-lin Lin, Lian-Qiang Feng, Yohei Doi, Nicole Stoesser, Guo-Bao Tian

**Affiliations:** Department of Immunology, Zhongshan School of Medicine, Sun Yat-sen University, 74 Zhongshan 2nd Road, Guangzhou 510080, China; Key Laboratory of Tropical Diseases Control (Sun Yat-sen University), Ministry of Education, Guangzhou 510080, China; Modernising Medical Microbiology, Nuffield Department of Medicine, University of Oxford, Oxford, United Kingdom; Department of Clinical Laboratory, Sun Yat-sen Memorial Hospital, Sun Yat-sen University, Guangzhou 510080, China; University of Pittsburgh Medical Centre, Pittsburgh, Pennsylvania 15261, USA

## Abstract

**Background:** *mcr-*1-mediated transmissible colistin resistance in Enterobacteriaceae is concerning, given colistin is frequently used as a treatment of last resort in multidrug-resistant Enterobacteriaceae infections. Reported rates of human *mcr-1* gastrointestinal carriage have historically been low.

**Objectives:** To identify trends in human gastrointestinal carriage of *mcr-1* positive and *mcr-1*-positive/cefotaxime-resistant Enterobacteriaceae in Guangzhou, China, 2011-2016, and investigate the genetic contexts of *mcr-1* in a subset of *mcr-1*-positive/cefotaxime-resistant strains using whole genome sequencing (WGS).

**Methods:** Of 8,022 faecal samples collected, 497 (6.2%) were *mcr-1*- positive, and 182 (2.3%) *mcr-1*-positive/cefotaxime-resistant. Trends in carriage were assessed using iterative sequential regression. A subset of *mcr-1*-positive isolates was sequenced (Illumina), and genetic contexts of *mcr-1* were characterised.

**Results:** We observed marked increases in *mcr-1* (now ~30% prevalence) and more recent (since January 2014) increases in *mcr-1*-positive/third-generation cephalosporin-resistant Enterobacteriaceae human colonisation (p<0.001). Sub-cultured *mcr-1*-positive/third-generation cephalosporin-resistant isolates were commonly multi-drug resistant.

WGS of 50 *mcr-1*/third-generation cephalosporin-resistant isolates (49 *Escherichia coli*; 1 *Klebsiella pneumoniae*) demonstrated bacterial strain diversity (39 *E. coli* sequence types); *mcr-1* in association with common plasmid backbones (IncI, IncHI2/HI2A, IncX4) and sometimes in multiple plasmids; frequent *mcr-1* chromosomal integration; and loss of the *mcr-1*-associated insertion sequence IS*Apl1* in some plasmids. Significant sequence similarity with published *mcr-1* plasmid sequences was consistent with spread amongst pig, chicken and human reservoirs.

**Conclusions:** The high positivity rate (~10%) of *mcr-1* in multidrug-resistant *E. coli* colonising humans is a clinical threat; the diverse genetic mechanisms (strains/plasmids/insertion sequences) associated with *mcr-1* have likely contributed to its dissemination, and will facilitate its persistence.

## INTRODUCTION

The polymyxins, including colistin, are antibiotics of last resort for managing multidrug-resistant Gram-negative infections, including *Pseudomonas aeruginosa, Acinetobacter* spp. and Enterobacteriaceae. Acquired resistance in these organisms has historically largely been due to: mutation-based modifications to cell wall lipopolysaccharides; utilization of efflux pumps; and capsule formation[1]. Transmissible colistin resistance, encoded by *mcr-1*, was recently identified in *Escherichia coli* and *Klebsiella pneumoniae* isolates from hospitalized humans, animals (pigs) and raw meat (pigs and chicken) in China[2], with higher rates in animal samples (~19% versus ~1% in humans).

Subsequently, *mcr-1*-harboring strains have been variably identified in humans, animals and raw meat in Europe, Asia, North America, South America and Africa (e.g.[3–13]. These strains have predominantly been *E. coli*[14] or *Salmonella* spp.[3, 5], with up to 20% carriage prevalence in swine and poultry[6, 11], and *mcr*-1-positive isolates from chickens as early as the 1980s in China and 2007 in France[6, 15]. Prevalence in humans remains low[2, 4, 7, 10], and mostly restricted to hospitalized patients[16]. However *mcr-1* can be carried in the healthy human gut[17].

The association of *mcr-1* with other broad-spectrum resistance mechanisms, such as extended-spectrum β-lactamases (ESBLs) and/or carbapenemases[18–23], has major clinical implications. As polymyxins are last-line therapies for carbapenem-resistant infections, widespread, joint dissemination of these mechanisms would represent a major problem for case management. The identification of *mcr-1* in multiple plasmid types, including IncI2[2, 23], IncHI2[20], IncX4[12, 23], IncP[24] and IncF[21] replicons, is consistent with multiple *mcr-1* mobilization events, potentially facilitating the association of *mcr-1* with other resistance mechanisms, thereby creating multidrug-resistant bacterial hosts.

Here we investigate human faecal carriage of *mcr-1* in Guangzhou, China, over 5 years, using whole-genome sequencing (WGS) to characterize the genetic structures involved in a subset of isolates and identify associations with other antimicrobial resistance genes.

## MATERIALS AND METHODS

8,022 consecutive faecal samples, initially collected to detect ESBL-producing Enterobacteriaceae, were submitted to the laboratory at Sun-Yat Sen Memorial Hospital 1^st^ April 2011-31^st^ March 2016. Samples came from three hospitals (Sun-Yat Sen Memorial Hospital, Guangdong General Hospital, and Qingyuan People’s Hospital) in Guangzhou, the capital city of Guangdong province, serving a population of ~15 million over ~10,000 km^2^ in southeast China. All samples were obtained continuously, except for January 2012, February 2013 and February 2014 (holiday months of the Spring Festival; staff shortages). Ethical approval for the study was given by Sun Yat-Sen University; individual consent for the use of faecal samples was obtained for all patients.

Faecal samples were plated onto Columbia blood agar (CBA) without antibiotic within 2 hours of collection and incubated for 18-24 hours at 37°C. Subsequently, up to 10 colonies with the appearance of Enterobacteriaceae were sub-cultured to MacConkey agar (Thermo Fisher, Beijing, China) supplemented with cefotaxime (2 mg/L), and species identification performed by 16S rDNA sequencing (Supplementary material). All cefotaxime-resistant Enterobacteriaceae isolates were stored in lysogeny broth (LB) with 30% glycerol at −80°C. Sweeps from the original CBA plates were also stored in LB with 30% glycerol at −80°C.

All frozen sweeps of cultured feces (n=8,022) and cefotaxime-resistant Enterobacteriaceae isolates (n=20,332) were subsequently re-cultured from frozen stock and screened for *mcr-1* by PCR (Supplementary material). Cefotaxime-resistant isolates were screened for *bla*_CTX-M_, and alleles determined by sequencing (Supplementary material). Species identification for *mcr-1* positive isolates was performed by API20E (bioMérieux, Marcy l’Etoile, France). Minimum inhibitory concentrations (MICs) were determined for all *mcr-1*-positive isolates by agar dilution and interpreted using the European Committee on Antimicrobial Susceptibility Testing (EUCAST) breakpoints (version 6.0), as well as the Clinical and Laboratory Standards Institute (document M100-S25).

All cefotaxime-resistant, *mcr-1*-positive Enterobacteriaceae of distinct species and with differing MIC profiles to May 2015 (n=45), and a random subset from June 2015-March 2016 (n=44/142 [31%]) were selected for WGS. Each stored, frozen isolate was inoculated into 50 mls LB, cultured overnight, centrifuged at 3000rpm, and transferred to the Beijing Genomics Institute (BGI) for DNA extraction and WGS. DNA was extracted using the cetyl-trimethyl-ammonium bromide (CTAB)/chloroform method (Supplementary material). Multi-locus sequence type was available for a subset (25/187) using a standard approach (protocols at http://mlst.warwick.ac.uk/mlst/dbs/Ecoli).

DNA extracts were sequenced on the Illumina HiSeq 4000 platform, using both paired-end (150bp reads, ~350bp insert) and mate-pair (50bp reads, ~6kb insert) approaches (n=69 isolates) or paired-end reads only (n=20 isolates; Table S1). Libraries were prepared using standardized protocols incorporating fragmentation by ultra-sonication, end repair, adaptor ligation, and PCR amplification (Supplementary material).

Preliminary species identification for isolates was derived from WGS using Kraken[25]; read-data were then mapped to a species-specific reference[26]. Hybrid *de novo* assemblies of paired-end and mate-pair data, or paired-end data alone, were generated using SPAdes[27] version 3.6 with the “--careful" option, a set of automatically determined k-mer values (21, 33, 55, 77), and by removing contigs <500bp or with k-mer coverage <1. *In silico* MLST was determined using BLASTn and publicly available databases for relevant species (http://mlst.warwick.ac.uk/mlst/dbs/Ecoli, http://bigsdb.web.pasteur.fr/klebsiella/klebsiella.html). Resistance genes were identified from *de novo* assemblies using an in-house, curated database of resistance genes[28] and BLASTn/mapping-based identification (scripts available at: https://github.com/hangphan/resisType). Contigs were annotated using PROKKA[29]. Contigs containing *mcr-1* were defined as “plasmid” if they contained annotations consistent with plasmid-associated loci and no obvious chromosomal loci, or “chromosomal” if >75% of annotations (excluding hypothetical proteins) were consistent with a chromosomal location.

The integrity of assemblies containing *mcr-1* in chromosomal locations was assessed using the REAPR tool[30]. We excluded any sequences with assembly sizes ≥6.5Mb; mixed calls from *in silico* MLST or species identification; and mismatches between available laboratory-based and WGS *in silico* MLST.

Phylogenies were reconstructed using IQtree[31] with a generalized time reversible nucleotide substitution model and a gamma distribution allowing for substitution rate variation among sites (GTR+G setting), using a starting tree inferred by maximum parsimony. Branch lengths were corrected for recombination using ClonalFrameML (default settings, kappa estimated from IQtree=2.75543)[32]. Insertion sequences (IS) were downloaded from ISFinder (https://www-is.biotoul.fr); sequence assemblies were queried with BLASTn (presence requiring >95% sequence identity over >90% of the reference sequence length) against this database.

Circularization of *mcr-1*-harboring plasmid contigs was confirmed using Bandage[33]. For single *mcr-1*-harboring plasmid contigs which could not be fully circularized, we also used Bandage to visualize the sequencing assembly graph generated by SPAdes and manually resolved the most likely *mcr-1* plasmid structures based on node (contig) linkage and contig coverage (Supplementary material).

Iterative sequential regression (ISR) in R was used to characterize trends in faecal *mcr-1* positivity (Supplementary material). The phylogeny was represented in the interactive tree of life viewer (iTOL v3, http://itol.embl.de).

Links to raw sequencing reads for the isolates included in analyses are available through the NCBI BioProject accession number: PRJNA354216 (Table S1).

## RESULTS

Of 8,022 samples collected during the 5-year study, 497 (6.2%) were *mcr-1*- positive, and 182 (2.3%) *mcr-1*- positive and cefotaxime-resistant (Fig. S1). The proportion of both *mcr-1*-positive and *mcr-1*-positive/cefotaxime-resistant faecal samples increased significantly during the study (p<0.001). For *mcr-1*-positive/cefotaxime-resistant samples, this was driven by a specific increase after January 2014 (p<0.001, Fig.1; 95% CI for estimated date of trend change: 01/April/2013-01/Nov/2014).

**Figure 1.**
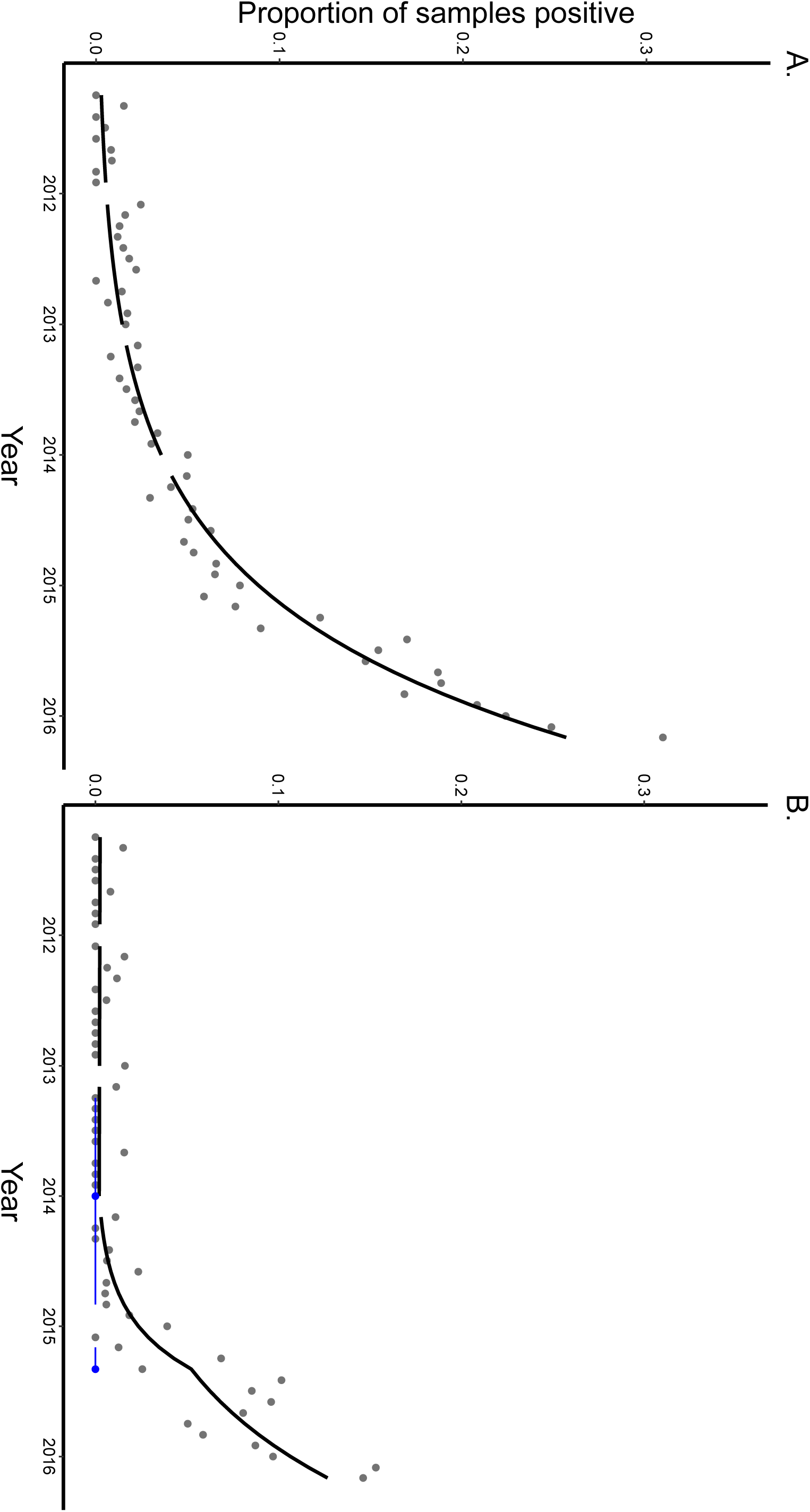
Monthly proportions of *mcr-1*-positive human faecal samples in Guangzhou, China, 2011-2016 for: Panel A, faecal samples harbouring *mcr-1*-positive isolates, and Panel B, faecal samples harbouring *mcr-1*-positive/cefotaxime-resistant isolates. Black line represents estimated prevalence by iterative sequential regression (ISR), with gaps representing months with missing data. Blue lines at the base of the graph represent 95% confidence intervals around the breakpoints estimated by the ISR model.

From faecal samples harbouring *mcr-1*-postive/cefotaxime-resistant Enterobacteriaceae, 187 distinct Enterobacteriaceae isolates from 179 individuals were identified (*E. coli*, n=173; *K. pneumoniae*, n=13; *Enterobacter cloacae*, n=1). 90 (50%) of these 179 individuals were male, median age 52 years (inter-quartile range [IQR: 39-69 years). 144/179 (80%) individuals were hospital in-patients at the time of sampling (median length of stay 14 days [range: 3-258 days; IQR: 7-21 days]).

The 89 sequenced isolates comprised *E. coli* (n=79), *K. pneumoniae* (n=9), *Enterobactercloacae* (n=1), based on 16S rDNA sequencing. Of these, however, 30/45 consecutive isolates pre-May 2015 and 9/44 randomly selected isolates post-June 2015 failed sequencing quality control metrics (Table S1), leaving 50 sequences for analysis (49 *E.coli*, 1 *K. pneumoniae*). For *E. coli*, these represented 39 sequence types (STs)(Fig.2), ST156 being the most common (n=4), with most isolates (n=33) representing singleton STs. Other global disease-causing lineages were also identified, including ST131, ST155 and ST405[34], and for *K. pneumoniae*, ST15. One pair of isolates was separated by 0 SNVs (SYSU0077/SYSU0078, 7 days apart), and one pair by 3 SNVs (SYSU0041/SYSU0049, 28 days apart), representing likely direct/indirect transmissions between patients 146 and 150, and 95 and 104, respectively. All other strains were >1126 SNVs apart.

**Figure 2.**
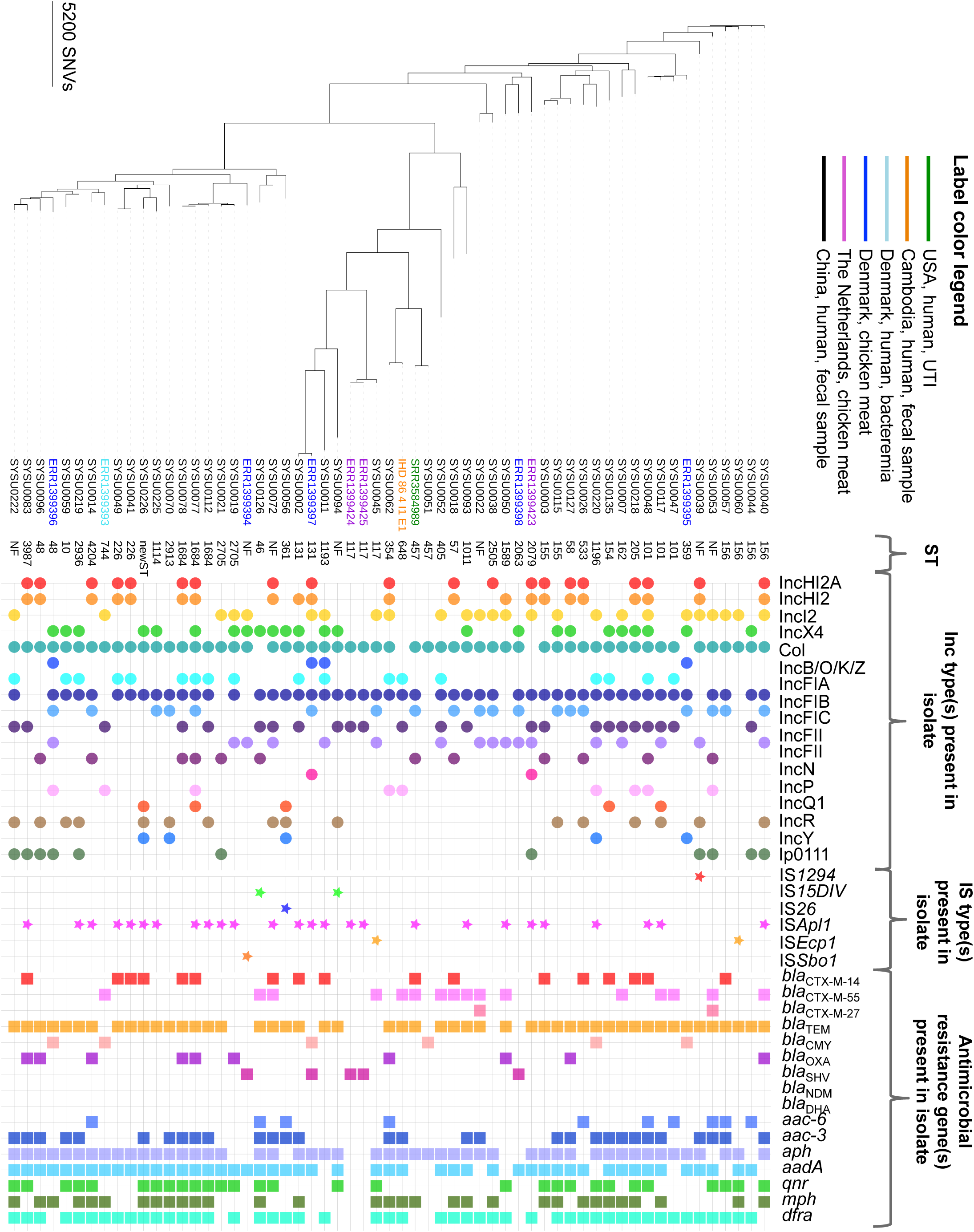
Phylogeny of sequenced *Escherichia coli* study isolates, plus available reference sequences from NCBI (n=11), and associated sequence types (ST; “NF” denotes “not found”); plasmid incompatibility groups; insertion sequences within 5kb of *mcr-1* on either the same contig or contigs associated through the assembly graph; and antimicrobial resistance genes present within the isolates (presence represented by respective coloured shapes).

A novel *mcr-1* gene variant was identified with a G3A mutation, leading to loss of the first methionine in the translated protein (n=1; SYSU0052). A separate isolate had *mcr-1* disrupted by an IS*1294* element, previously described downstream of *mcr-1* (n=1; SYSU0039)[8]. The former did not affect colistin resistance phenotype (MIC colistin=8 mg/L, polymixin B=16 mg/L); the latter, as expected, was colistin-susceptible (MIC colistin<0.25 mg/L, polymixin B=1 mg/L).

128/187 (68%) isolates contained *bla*_CTX-M_ by PCR/sequencing (Table S1). Rates of co-resistance across antimicrobial classes in *mcr-1*-positive/cefotaxime-resistant isolates were high (Table 1), with only carbapenems, nitrofurantoin and tigecycline demonstrating susceptibility rates ≥80%. Resistance was similar in sequenced isolates (Table 1). Of these, 40/50 (80%) harboured *bla*_CTX-M_, including: (i) group 9-like alleles: 14(n=15), 27(2), 65(5), 110(1), 130(1); (ii) group 1-like alleles: 1-like(n=1), 11(1), 55/55-like(16), 136(1); and (iii) hybrid alleles: 64(2), 123(2). Eight (16%) isolates had 2 *bla*_CTX-M_ variants. In three isolates (carrying *bla*_CTX-M-14_,_27/55,55/123_), a *bla*_CTX-M_ variant was located on the same plasmid contig as *mcr-1*. The presence of *bla*_CMY_(2, 42-like [N90T]) variants explained cefotaxime resistance in two isolates. For seven (14%) isolates, no cefotaxime resistance mechanism could be identified (Table S1). Multiple other resistance genes identified included: (i) *bla*_TEM-1_ (n=37), *bla*_TEM-135_(2), *bla*_TEM-116_(1), *bla*_TEM-181_(1), *bla*_TEM-191-like_(1[K227E]), *bla*_TEM-30-like_(1[S241R, A148P]); (ii) *bla*_OXA-1_(2) and/or *bla*_OXA-10_(8), *bla*_OXA-10-like_(1[truncated, S14*]); (iii) *fosA3* (n=23); (iv) *qepA* (n=1) or *qnrS1* (n=12); *rmtB*(2) or *armA*(1); and (v) *aac(6’)-Ib-cr* (6) (Table S1).

**Table 1.**
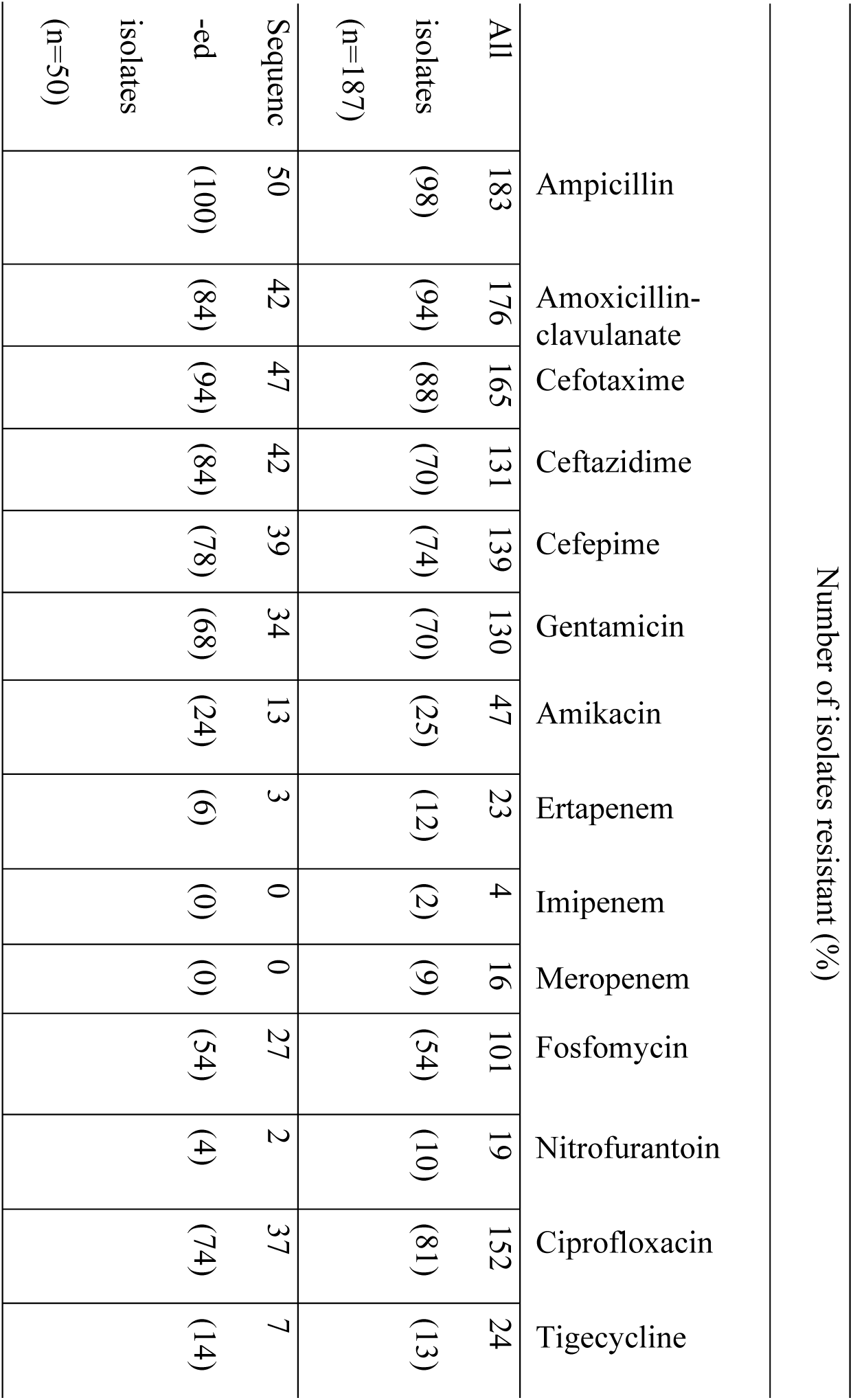
**Resistance profiles of *mcr-1*-positive/cefotaxime-resistant study isolates.**

The genetic context of *mcr-1* was similarly diverse, supporting the potential for *mcr-1* to have spread widely, and in combination with other resistance genes. In 41/50 (82%) isolates *mcr-1* was located on plasmid-associated contigs, in 4 (8%) on chromosomal contigs, in 2 (4%) likely on chromosomal contigs, and in 3 (6%) the location was unclear due to assembly/annotation limitations. *mcr-1* copy number was estimated at up to 3.54 (median 0.97, inter-quartile range: 0.79-1.39); likely due to its presence either on multi-copy plasmids; as multiple copies within the same plasmid structure (SYSU0077 [IncH]; SYSU0093 [IncI2]); or on different plasmids within the same isolate (SYSU0072 [IncH and IncX4]; SYSU0220 [novel plasmid described below, IncI2-like plasmid backbone]). Insertion sequences may contribute to plasmid-plasmid and plasmid-chromosome rearrangements and gene mobilization. We found 99 different IS types in the 50 sequenced isolates, with median 19.5 (range: 9-30) IS types per isolate. The commonest were IS*3* (46/50 isolates), IS*186B*, IS*2* (42/50 each), and IS*150* (39/50). IS*Apl1*, most frequently associated with *mcr-1* mobilization to date, was found in 21/50 (42%) isolates.

In 19/41 (46%) plasmid-associated isolates *mcr-1* was co-located on the same contig as an IncI2 replicon (n=17), or plausibly on an IncI2 plasmid based on assembly visualization (n=2). Eleven of these 19 structures could be circularized, likely representing complete plasmid structures; this included the earliest sequenced plasmid in this study, and the earliest sequenced *mcr-1* plasmid, to our knowledge (isolated May 2011). In these 11 plasmids, the backbone, *mcr-1* location and wider gene synteny were largely preserved, consistent with ancestral acquisition of *mcr-1* and subsequent evolution of the plasmid structure by mutation/recombination (e.g. *repA* and the transfer region, Fig.3) and acquisition/loss of smaller mobile genetic elements (MGEs). The last included variable presence of IS*Sen6*, a *Salmonella* spp.-associated MGE (SYSU0011, SYSU0050, SYS0093); IS*Ecp1*-associated acquisition of *bla*_CTX-M-55_ (SYSU0060, SYSU0045); and variability in the shufflon region adjacent to the *rci* gene in the C-terminal region of the *pilV* gene).

**Figure 3.**
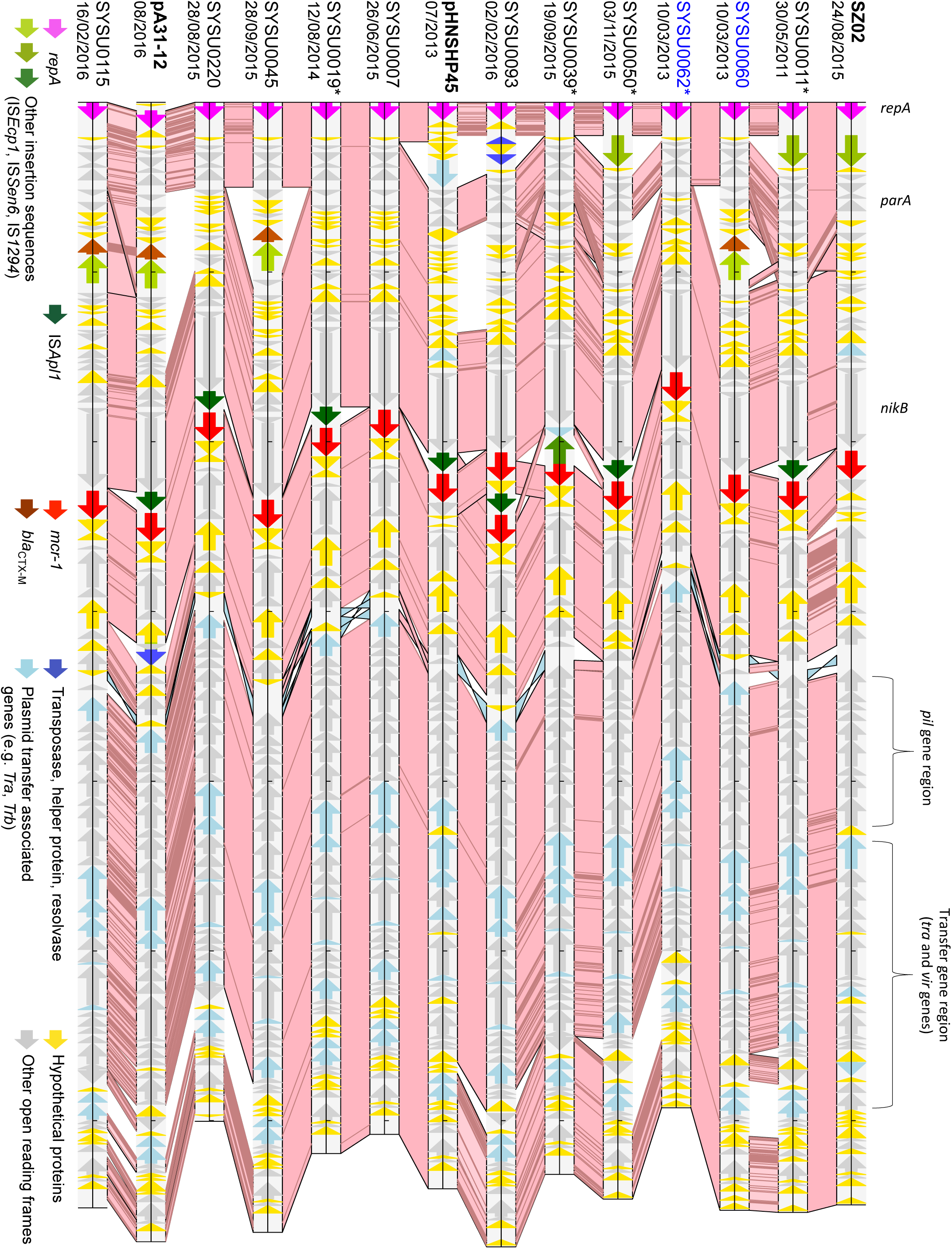
Alignment of *mcr-1* IncI2 study plasmid contigs (incomplete, denoted by *) and circularized plasmid sequences, and three reference plasmid sequences (sequence labels in bold followed by dates of host strain isolation). Pink/blue cross-links between aligned sequences demonstrate regions of sequence homology (BLASTn matches of ≥500bp length, >95% sequence identity); dark regions within these cross-links demonstrate sequence variation. Tick marks represent 10kb of sequence. Sequences are ordered based on relatedness inferred from multiple sequence alignment (Kalign-EBI). Coloured arrows demonstrate ORFs of particular relevance. Two plasmids were obtained from isolates cultured from the same patient (sequence labels in blue).

These 11 IncI2 plasmids were also genetically highly similar to three previously described plasmids: SZ02 (Accession number: KU761326 [Fig.3], isolated from an outpatient’s blood culture, August 2015, Suzhou, China, ~1,400km away[23]); the *bla*_CTX-M-55_-harboring pA31-12 (Accession number: KX034083.1, chicken, August 2012, Guangzhou), and pHNSHP45 (Accession number: KP347127.1 [Fig.3], pig, July 2013, Shanghai), consistent with the dissemination of this plasmid in human cases/carriers, pigs and chickens across China.

In relation to *mcr-1*, we observed variable presence of IS*Apl1* (n=5/11 [45%] study isolates, reference plasmids: pHNSHP45, pA31-2). When IS*Apl1* was present, however, it was consistently located 186bp upstream of *mcr-1* between *mcr-1* and *nikB*. The most likely hypothesis is a historical introduction of IS*Apl1* and *mcr-1* into the IncI plasmid backbone, with subsequent loss of IS*Apl1*. We also observed highly genetically related plasmid backbones in two different *E. coli* host strains (ST156, 354; SYSU0060, SYSU0062) isolated from the same patient on the same day, potentially consistent with within-host transfer, as well as highly genetically related plasmids within different *E. coli* host strains and human individuals across periods of time consistent with direct transmission/acquisition from common sources (SYSU0019, SYSU0007).

In 14 (34%) isolates *mcr-1* co-localized to a contig with either IncHI2A or IncHI2+HI2A (n=8), or was plausibly associated with an IncHI2 plasmid based on assembly visualization. These plasmids are typically large, and we were unable to reconstruct any complete IncHI2A plasmids from the sequencing data. Four contigs were short (<3,000bp); the rest are represented in Fig.4, alongside two closed reference *mcr-1* IncHI2 plasmids, pSA26 (Accession number: KU743384, bacteraemia, Saudi Arabia), and pHNSHP45-2 (Accession number: KU341381, pig faeces isolate, Shanghai, China). This alignment demonstrates a relatively homogenous plasmid backbone circulating within the study population between 2014-early 2016, likely derived from an ancestral plasmid structure similar to the reference plasmids, with some rearrangement/inversion events, particularly around a region enriched for resistance genes and transposable elements (Fig.4). Within the study isolates, sampled over 3.5 years, we observed apparently frequent IS/transposon-associated indel events, including of IS*Apl1*, which was either at the 5’ end of *mcr-1* (e.g. SYSU0003), flanking it on both sides (e.g. SYSU0014), or absent (e.g. SYSU0026). As *mcr-1* was located in the same wider genetic context in 9/10 study plasmid contigs, this likely represents a single IS*Apl1*-*mcr-1*-IS*Apl1* acquisition and subsequent loss of IS*Apl1* elements[35]. Alternatively it could represent a “hotspot” for multiple insertion events of this complex. We also observed a *mcr-1* duplication event within an otherwise identical contiguous plasmid sequence in two isolates taken 7 days apart from different patients (both *E. coli* ST1684; SYSU0077, SYSU0s078); and an inversion of IS*Apl1*-*mcr-1* within the plasmid backbone (SYSU0002), all consistent with high rates of plasticity involving the IS*Apl1*-*mcr-1* and IncHI2/HI2A structures. Other variation in the IncHI2+HI2A backbone in study isolates included: (i) single nucleotide variants (SNVs), (ii) small indels (<20bp), (iii) clustered SNVs suggestive of local recombination events, and (iv) gain and loss of several insertion sequences.

**Figure 4.**
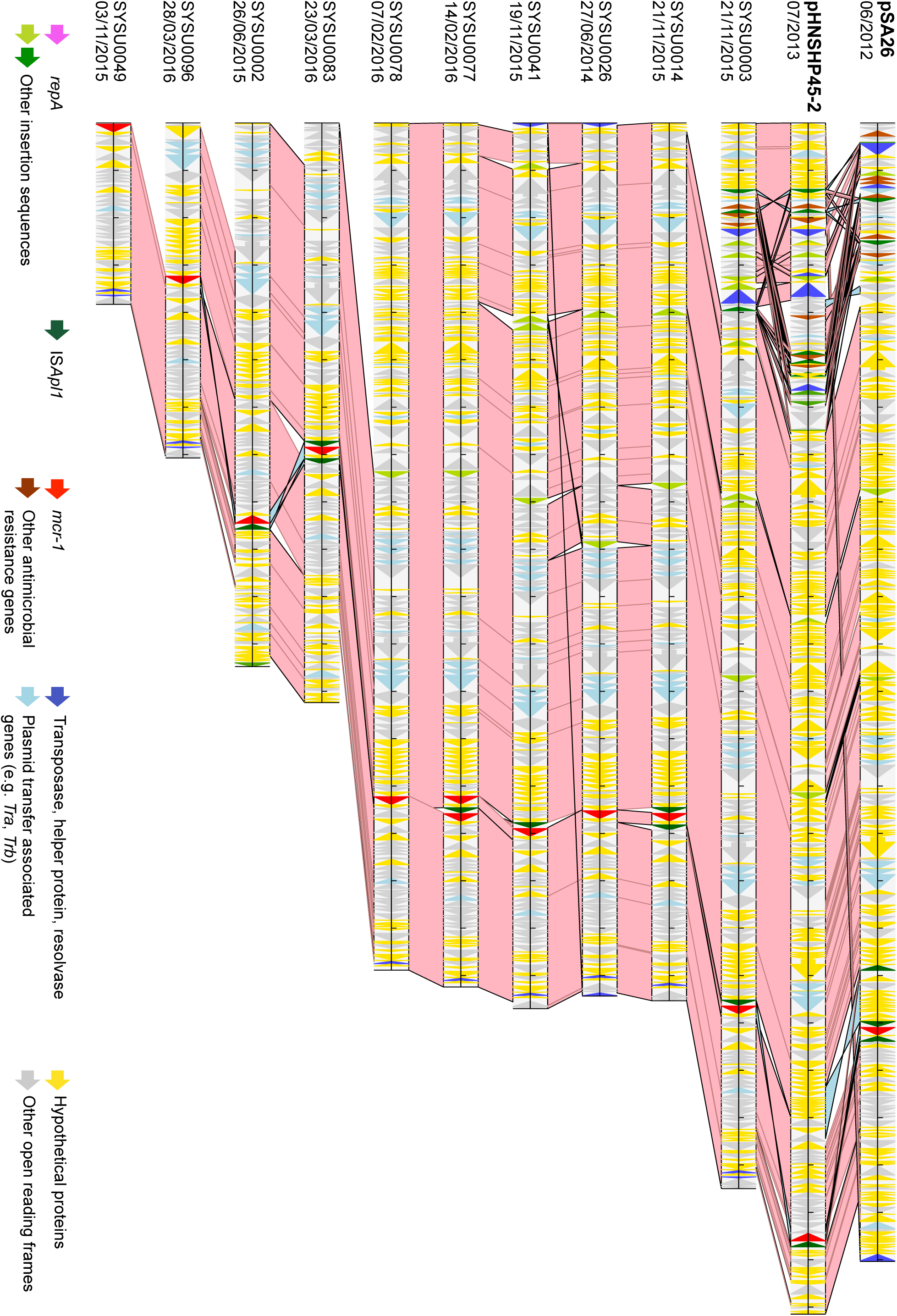
Alignment of *mcr-1* IncHI2/HI2A plasmid contigs and two reference plasmid sequences (sequence labels in bold followed by dates of host strain isolation). Pink/blue cross-links between aligned sequences demonstrate regions of sequence homology (BLASTn matches of ≥500bp length, >95% sequence identity); dark regions within these cross-links demonstrate sequence variation. Tick marks represent 10kb of sequence. Coloured arrows demonstrate ORFs of particular relevance.

In seven (15%) isolates *mcr-1* co-localized to (n=6) or was plausibly associated with (n=1) an X4-harboring contig; in four the contigs could be fully circularized, likely representing complete plasmid structures (Fig.5). These sequences had a highly conserved, syntenic plasmid backbone, with limited nucleotide and indel variation, including to two reference IncX4-*mcr-1* plasmids, SZ04 (inpatient peritoneal fluid, Suzhou, China) and pAF48 (urine, Johannesburg, South Africa). IS*Apl1* was absent in all of these plasmids; the only consistently associated IS was an IS*26* or highly similar IS*15*DIV element 4.7kb upstream of *mcr-1*.

**Figure 5.**
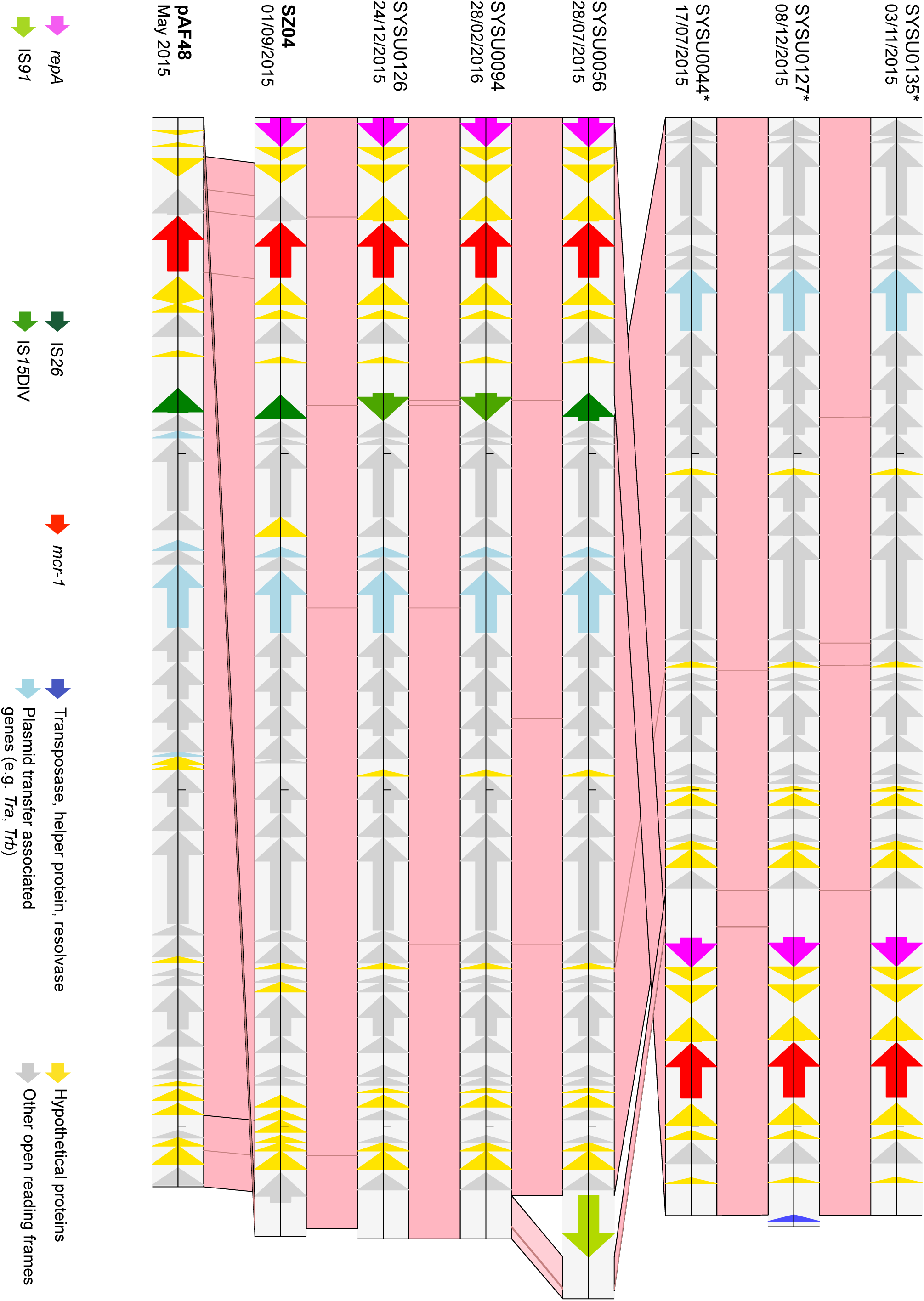
Alignment of *mcr-1* IncX4 study plasmid contigs (incomplete, denoted by *) and circularized plasmid sequences, and two reference plasmid sequences (sequence labels in bold followed by dates of host strain isolation; the top three sequences are incomplete plasmid contigs). Pink/blue cross-links between aligned sequences demonstrate regions of sequence homology (BLASTn matches of ≥500bp length, >95% sequence identity); dark regions within these cross-links demonstrate sequence variation. Tick marks represent 10kb of sequence. Coloured arrows demonstrate ORFs of particular relevance.

In 11 (27%) isolates no co-located or plausibly associated plasmid replicon could be identified, and in one isolate (16EY645) we identified a new *mcr-1* harbouring plasmid structure, p16EY645_1, of 48,944bp (Fig. S2). 94% of p16EY645_1 shared 99% sequence identity with a larger IncP plasmid pHNFP671 (82,807bp, accession number: KP324830.1), isolated from pig faeces obtained prior to 2014 in Guangzhou, China (the study region); but p16EY645_1 itself has no typeable replicon.

Chromosomal *mcr-1* integration was confirmed in four *E. coli* STs (457, 1114, 1684, 2936), and plausibly occurred in two others (101, 2705), consistent with at least four-six independent chromosomal integration events in diverse *E. coli* strains (12% sequenced *E. coli* isolates), providing capacity for widespread dissemination at the organism level. In three of these isolates (SYSU0021, SYSU0219, SYSU0112), IS*Apl1* flanked *mcr-1* on both sides (with AT-AT, AC-CC as the composite transposon target site sequences [TSS], or indeterminate TSS due to contig breaks). In one isolate *mcr-1* was downstream of a single IS*Apl1* unit (SYSU0225); in one flanked by a previously un-described IS*Ec23*/IS*30*-like/IS*1294* structure (SYSU0051); and in one case inserted within a much larger integrative element harbouring the *iee* gene encoding for an IS excision enhancer, which promotes excision of members of the IS*3*, IS*1*, and IS*30* families, including IS*Apl1*[36](SYSU0047).

## DISCUSSION

Widespread transmissible colistin resistance is of major concern in the current era of multidrug-resistant Gram-negative infections, as colistin is commonly used to treat infections caused by these organisms, despite its nephrotoxicity, and issues with determining susceptibility and appropriate dosing regimens[37, 38]. In this, the largest study of faecal carriage and WGS of *mcr-1* isolates to date, we observed alarming sequential increases in human faecal carriage of *mcr-1*-positive and cefotaxime-resistant, third-generation cephalosporin-resistant *E. coli* over 5 years. We also found high genetic plasticity and multidrug resistance of MGEs harbouring *mcr-1* in these regional data from Guangzhou, China, that may explain some of the dramatic recent increases in human faecal carriage over the last two years.

IncI plasmids are narrow host-range plasmids[39], commonly isolated from *E. coli* and *Salmonella* spp., and implicated in the spread of the ESBL gene *bla*_CTX-M_ amongst Enterobacteriaceae, particularly in China[40, 41]. CTX-M-55 has been found in *E. coli* in animals in China, and CTX-M-55/55-like variants were seen with *mcr-1* in IncI2 plasmids in two different *E. coli* strains (ST156, ST117) in this study. Previously, it has been postulated that an IncI2 plasmid backbone acquired a 3,080bp IS*Ecp1*-*bla*_CTX-M-15_ complex (GAAAA TSS) from an IncA/C plasmid, with a mutation resulting in conversion to *bla*_CTX-M-55_ (V77A) within this structure[42]. The same signature sequence was observed in our isolates, as well as another recently sequenced IncI2 plasmid harbouring *bla*_CTX-M-55_ from a chicken *E. coli* isolate in Guangzhou, China (August 2012)[43], consistent with the spread of this plasmid across humans and several animal species. *bla*_CTX-M-64_ was also observed on an IncI *mcr-1* plasmid in this study (SYSU0115, ST155); this allele has previously been postulated to have arisen from recombination between *bla*_CTX-M-14_ and *bla*_CTX-M-15_ on IncI plasmid backbones within food animals in China[42].

IncHI2/HI2A plasmids are typically large (>250kb)[44], multidrug-resistant plasmids that have been associated with a range of antibiotic and metal resistance genes in *Salmonella* spp. and *E. coli* isolated from humans and food-producing animals[45]. Similar to the IncI plasmids, our genetic analyses suggest an initial *mcr-1* acquisition event, in this case by direct transposition of a composite IS*Apl1* transposon, and subsequent loss of IS*Apl1* units and rearrangements of the plasmid backbone.

In at least four (8%) sequenced isolates, *mcr-1* had been integrated into the chromosome, a phenomenon observed in at least two other studies (*E. coli* ST156, Beijing; IS*Apl1* composite transposon, TG-TG TSS[46]; *E. coli* ST410, Germany; IS*Apl1* signatures consistent with composite transposition and then loss of downstream IS*Apl1*, CA-CA TSS[47]). Our data suggest two additional *mcr-1* chromosomal integration events - one in association with an integrative element promoting IS activity, and one in association with IS*Ec23*/IS*30*-like/IS*1294* element, which has likely played a role in the mobilization of *bla*_CMY-2_ in *Salmonella* spp. and *E. coli*[48]. The multifarious means by which chromosomal integration of *mcr-1* appears to occur is of major concern, as its presence in the chromosome may facilitate more stable inheritance of this gene within multiple *E.coli* strains.

IS*Apl1* is an IS*30*-like element derived from *Actinobacillus pleuropneumoniae*, a Gram-negative organism of the Pasteurellaceae family, known to cause pleuropneumonia in pigs[49], a highly infectious disease frequently warranting herd-based antimicrobial treatment. IS*30*-like elements form circular intermediates during transposition, and have been shown *in vitro* to result in 2bp target site repeats (TG, TT, AA, GG) on direct transposition[49]. The variable extent of diversity observed within the plasmid families here (IncHI2/HI2A and IncI > IncX4) may reflect different evolutionary rates for these plasmid families, or may be consistent with earlier acquisition of IS*Apl1*-*mcr-1* composite transposons by IncHI2/HI2A and IncI2, most likely within poultry and pig farms in which regular colistin exposure represents a major selection pressure[2]. The acquisition of *mcr-1* by IncX4 plasmids appears to be unrelated to the presence of IS*Apl1*, and possibly associated with a rearrangement event involving IS*26/26*-like structures (e.g. IS*15*DIV).

This study has several limitations. Firstly, although we observed significant increases in faecal carriage of *mcr-1*-harboring isolates, we were unable to assess the possible risk factors associated with increasing incidence. Secondly, our sampling strategy targeted limited numbers of colistin-resistant isolates within faecal samples, and the diversity of *E.coli* and likely other Enterobacteriaceae strains in the gastrointestinal tract is substantial[50]. Our results could therefore underestimate the genetic diversity of strains and MGEs harbouring *mcr-1*, particularly within individual hosts. Finally, due to resource limitations we were unable to sequence all *mcr-1*-positive isolates, or to re-sequence those representing mixtures on sequencing. We were also unable to undertake any long-read sequencing, increasingly the method of choice in assembling plasmids.

Despite these limitations, we have demonstrated that human faecal carriage of *mcr-1* positive *E. coli* has increased dramatically in Guangzhou over the last two years, reaching similar proportions (20-30%) to animals (pigs, chickens) over the preceding 3-4 years in the same region of China[2]. Our genetic analyses suggest the rapid emergence of several major plasmid vectors of *mcr-1* within numerous multidrug-resistant *E. coli* strains carried by humans, and highlight the significant degree of plasticity in these plasmid vectors harbouring *mcr-1* over short periods of time.

## TRANSPARENCY DECLARATIONS

No conflicts of interest to declare.

## ACKNOWLEDGEMENTS

We would like to acknowledge the work of the Modernizing Medical Microbiology (MMM) sequencing pipeline team (Nicholas Sanderson) and informatics team (Ian Szwajca, Trien Do, Dona Foster) - collectively represented as the Modernizing Medical Microbiology Informatics Group.

## TRANSPARENCY DECLARATIONS

No conflicts of interest to declare.

## Supplementary tables/figures

**Table S1.** Summary details of study isolate source, phenotype, lab results and whole-genome sequencing metrics.

**Figure S1.** Monthly counts of: (i) all faecal samples collected, (ii) faecal samplesharbouring *mcr-1*-positive isolates, and (iii) faecal samples harbouring *mcr-1*-positive/cefotaxime-resistant isolates

**Figure S2.** Alignment of the novel, untypeable *mcr-1*plasmid pSYSU0220_1 and incPplasmid pHNFP671 (Accession number: KP324830).

